# Expanding the Zebrafish Genetic Code through Site-Specific Introduction of Azido-lysine, Bicyclononyne-lysine, and Diazirine-lysine

**DOI:** 10.1101/631432

**Authors:** Junetha Syed, Saravanan Palani, Scott T. Clarke, Zainab Asad, Andrew R. Bottrill, Alexandra M.E. Jones, Karuna Sampath, Mohan K Balasubramanian

## Abstract

Site-specific incorporation of un-natural amino acids (UNAA) is a powerful approach to engineer and understand protein function [1-4]. Site-specific incorporation of UNAAs is achieved through repurposing the amber codon (UAG) as a sense codon for the UNAA, a tRNA^CUA^ that base pairs with an UAG codon in the mRNA and an orthogonal amino-acyl tRNA synthetase (aaRS) that charges the tRNA^CUA^ with the UNAA [5, 6]. Here, we report expansion of the zebrafish genetic code to incorporate the UNAAs, Azido-lysine (AzK), bicyclononyne-lysine (BCNK), and Diazirine-lysine (AbK) into green fluorescent protein (GFP) and Glutathione-S-transferase (GST). We also present proteomic evidence for UNAA incorporation into GFP. Our work sets the stage for the use of UNAA mutagenesis to investigate and engineer protein function in zebrafish.

## Introduction

Site directed mutagenesis is a powerful approach to engineer and investigate protein function. In site directed mutagenesis one or more amino-acids in a protein is / are replaced with other canonical amino acid(s), often with properties in stark contrast with the original amino-acid (canonical is defined to encompass the 20 amino acids found in proteins) [7, 8]. The replacement of native amino-acids within a protein with other amino-acids allows functional dissection of individual amino-acid residues as well as domains in proteins. The ability to introduce UNAAs with novel properties in a site-specific manner is an important extension to site-directed mutagenesis and has the potential to revolutionize investigation and engineering of protein function. Site-specific incorporation of UNAAs has been made possible through the availability of UNAAs with a plethora of useful properties and the availability of repurposed codons and orthogonal tRNA and aminoacyl tRNA synthetases for these amino acids [9]. A large number of UNAAs are available which permit among others, photo-crosslinking, photo-uncaging, and click-chemistry.

Zebrafish is an attractive vertebrate model organism to uncover mechanisms of development and disease [10]. We set out to establish genetic code expansion strategies in zebrafish to facilitate structural and functional studies of proteins. Previous work has shown that azido-phenylalanine (Chen et al., 2017) and some lysine derivatives (Liu et al., 2017) could be introduced into zebrafish proteins through genetic code expansion. We sought to incorporate three useful amino-acids, Azido-Lysine, bicyclononyne-lysine, and Diazirine-lysine, which had not been previously incorporated into proteins in zebrafish. We show successful incorporation of these amino-acids into translation products of synthetic GFP and GST mRNAs injected into zebrafish embryos.

## Results and discussion

The strategy we employed to expand the genetic code in zebrafish embryos is shown in Figure 1A. Briefly, 1-2 cell zebrafish embryos were injected with a mixture carrying *in vitro* transcribed tRNA^CUA^ and mRNAs for the orthogonal aaRS (referred to the UNAA cocktail), UAG codon bearing mRNA for a target protein (GFP or GST) and an UNAA. The GFP used was fused to LifeAct, a short peptide that binds the F-actin cytoskeleton, so as to eliminate background autofluorescence, which might complicate evaluation of successful genetic code expansion [11]. Deiters and colleagues have successfully incorporated a variety of lysine derivatives, including a photo-cleavable amino acid, into zebrafish proteins [12]. In this work, we tested other lysine derivatives not used in their study, to further expand the tool kit for use in zebrafish. Diazirine-lysine and Azido-lysine generate reactive carbene and nitrene, respectively, upon exposure to ultraviolet light. The carbene and nitrene groups cause covalent cross-links with amino-acids in close proximity (∼3-12Å), allowing for investigation of direct binding partners [13, 14]. Azido-lysine can also be used in click chemistry reactions (strain promoted azide-alkyne cycloaddition-SPAAC. Bicyclononyne-lysine can be used in a different kind of click chemistry reactions (strain promoted inverse electron demand Diels-Alder cycloaddition-SPIDEAC) [15]. We used two different tRNA synthetases 1) *Methanosarcina mazei* PylRS (carrying the following amino acid substitutions Y360A and Y384F) for incorporating AzK and BCNK [16] and 2) *Methanosarcina barkeri PylRS (*L274M, C313A, Y349F) for integrating AbK [17]. In all cases, *Methanosarcina* Pyrrolysine tRNA^Pyl-CUA^ that base-pairs with UAG codon introduced into the mRNA was used, and is charged by either aaRS used in this work [18]. All GFP and GST mRNAs used contained an ochre UAA codon for translation termination.

**Figure 1:**
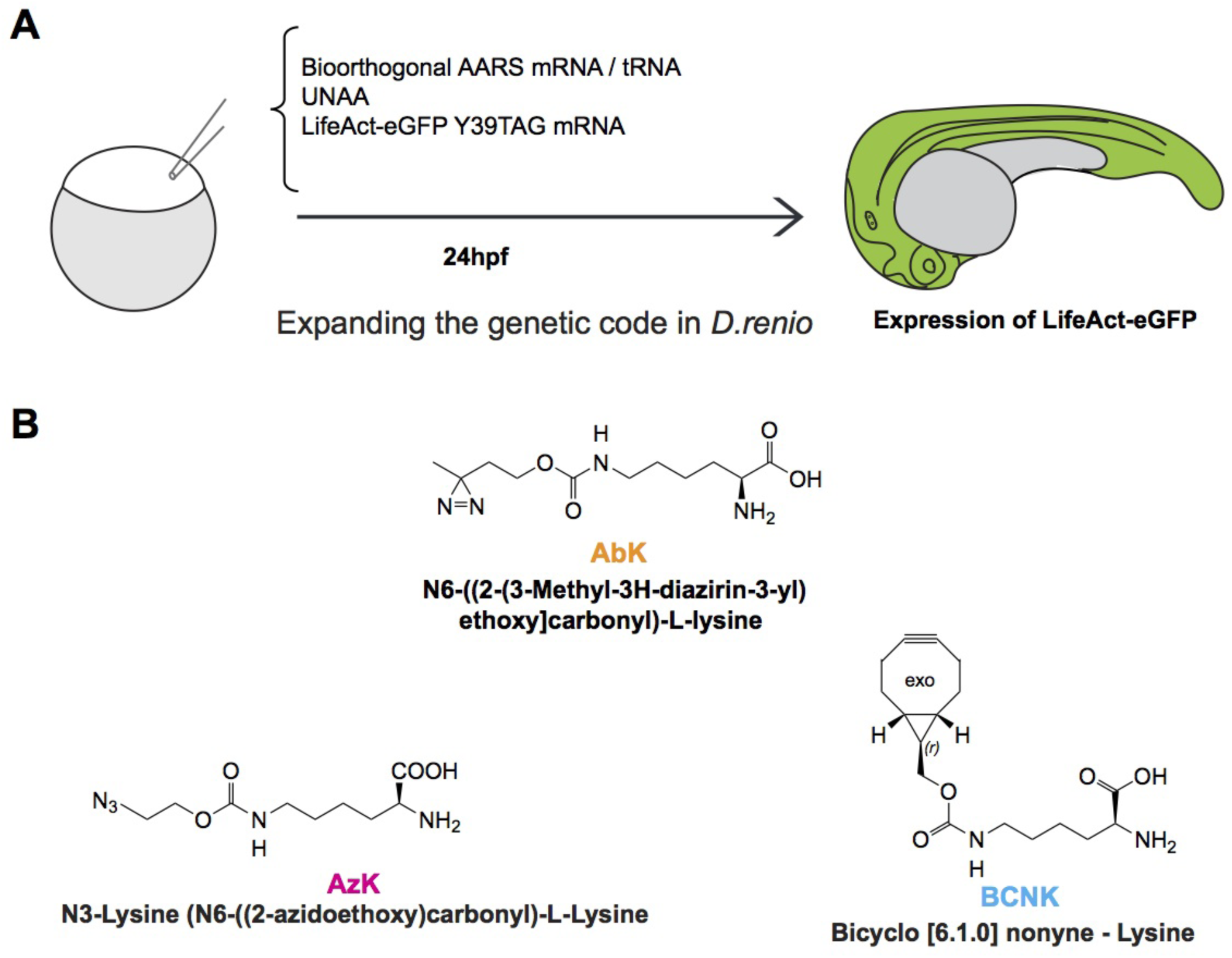
(A). Strategy for genetic code expansion in zebrafish embryos. (B). Structures of the UNAAs used in this study.

In embryos injected with LifeAct-eGFP mRNA, robust GFP expression was observed and at a subcellular level, as expected, the majority of LifeAct-eGFP localized to cell-cell junctions and in cortical speckles (Figure 2A and B). In control embryos where mRNA encoding LifeAct-eGFP Y39* (where a UAG codon replaced the codon for Y39) was injected together with the appropriate AzK/BCNK cocktail, but without the UNAA, no GFP signal was detected (Figure 2A). By contrast, when AzK/BCNK cocktail was injected along with the AzK, strong GFP expression was detected (Figure 2A). At the subcellular level, LifeAct-eGFP Y39AzK localized to cell-cell junctions and to cortical speckles, as observed in the case of LifeAct fused to wild-type eGFP. The fact that GFP expression was detected only in the presence of AzK strongly suggested the specificity of the AzK/BCNK cocktail in zebrafish embryos. To further validate site-specific incorporation of AzK into LifeAct-eGFP at position Y39, LifeAct-eGFP was immunoprecipitated using GFP-Trap and analysed by nano LC-ESI-MS/MS after digestion with trypsin. Proteomic studies further confirmed this suggestion. LifeAct-eGFP peptide (FSVSGEGEGDAT**<u>K</u>**GK) that incorporates azido-lysine at position 39 with a 113 Da adduct (the predicted mass of the added azido-moiety), was readily identified by mass spectrometry as a 2+ ion of 791.3606 m/z. We also successfully incorporated AzK into GST, and found that GST was detected through immunoprecipitation followed by western blotting in embryos injected with GSTF52* and AzK/BCNK cocktail with AzK, but not when AzK was not included (Figure 2D). Collectively, these experiments established that we had successfully expanded the genetic code of zebrafish to site-specifically introduce AzK.

**Figure 2:**
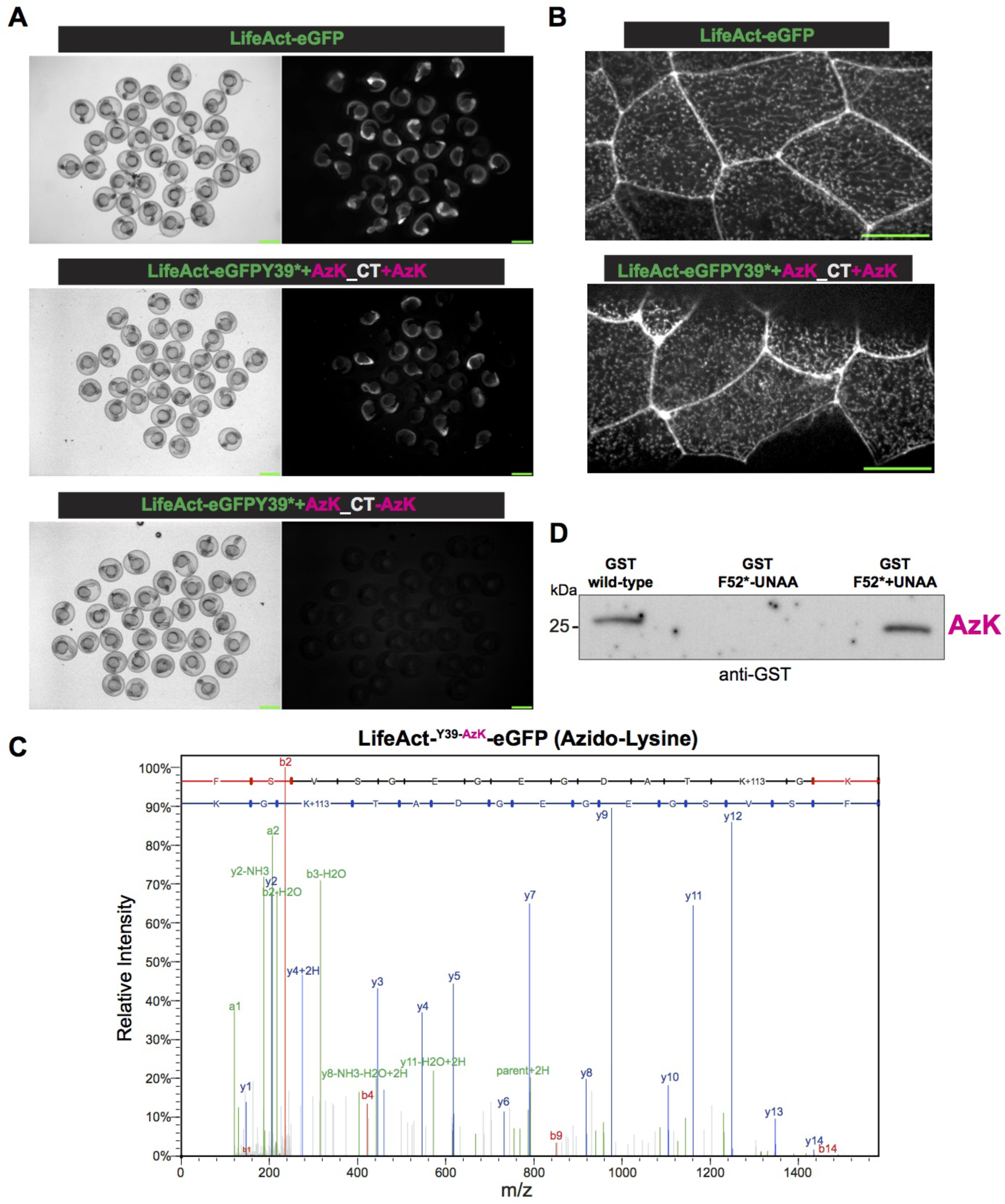
Site-specific incorporation of Azido-lysine in zebrafish embryos. (A). Shown are fields with multiple examples each of zebrafish embryos injected with mRNA(s) for LifeAct-eGFP (top panel) or LifeAct-eGFP Y39* (codon for Y39 converted to stop codon UAG) with the orthogonal tRNA^Pyl CUA^-mRNA aaRS for AzK/BCNK, with (middle panel) or without (bottom panel) AzK. Scale bar represents 1mm (B). Localization of LifeAct-eGFP or LifeAct-eGFP Y39AzK to cell-cell junctions and cortical speckles. Scale bar represents 20µm. (C). Mass-spectrometric identification of AzK in LifeAct-eGFP Y39AzK. Lysates were prepared from ∼1000 embryos injected with mRNA for LifeAct-eGFP Y39* with the orthogonal tRNA^Pyl-CUA^ -mRNA aaRS for AzK/BCNK with AzK. LifeAct-eGFP Y39AzK was immunoaffinity purified using GFP nanobodies and processed for mass spectrometry as described in methods. MS/MS spectrum of precursor m/z = 791.3606 (2+), corresponds to tryptic peptide FSVSGEGEGDAT**<u>K</u>**GK with Azido-Lysine incorporated at the indicated position. The full y-ion fragment series is highlighted in blue. (D). Shown is a IP-Western blot from zebrafish embryos injected with mRNA(s) for GST (left lane) or GFT F52* (codon for F52 converted to stop codon UAG) with the orthogonal tRNA^Pyl-CUA^ -aaRS pair for AzK/BCNK, without (middle lane) or with (right lane) AzK. Immunoprecipitations and western blots were performed with antibodies against GST.

We next tested for incorporation of BCNK into proteins in zebrafish embryos using the strategy employed in Figure 2, except that BCNK was used instead of AzK, since the same aaRS is capable of charging the tRNA with AzK and BCNK. We found GFP expression when embryos were injected with BCNK (Figure 3A; BCNK panel and see Figure 3B for localization to cell-cell junctions and cortical speckles), but not in the absence of BCNK (Figure 2A). Similarly, GST expression could be readily detected when mRNA for GST F52* was coinjected with AzK/BCNK cocktail and BCNK (Figure 3C). We detected a faint band in the sample that was not injected with BCNK. It is possible that there may be a very low level / inefficient charging of tRNAPyl-CUA with a cellular aaRS or an inefficient binding of a natural amino acid to aaRS AzK/BCNK. Finally, we tested if AbK could be introduced into proteins in zebrafish through genetic code expansion. To this end, we injected zebrafish embryos with AbK cocktail with or without AbK. In these experiments, we found that GFP expression and fluorescence (Figure 3A and B) and GST expression were detected only when embryos were injected with AbK (Figure 3C).

**Figure 3:**
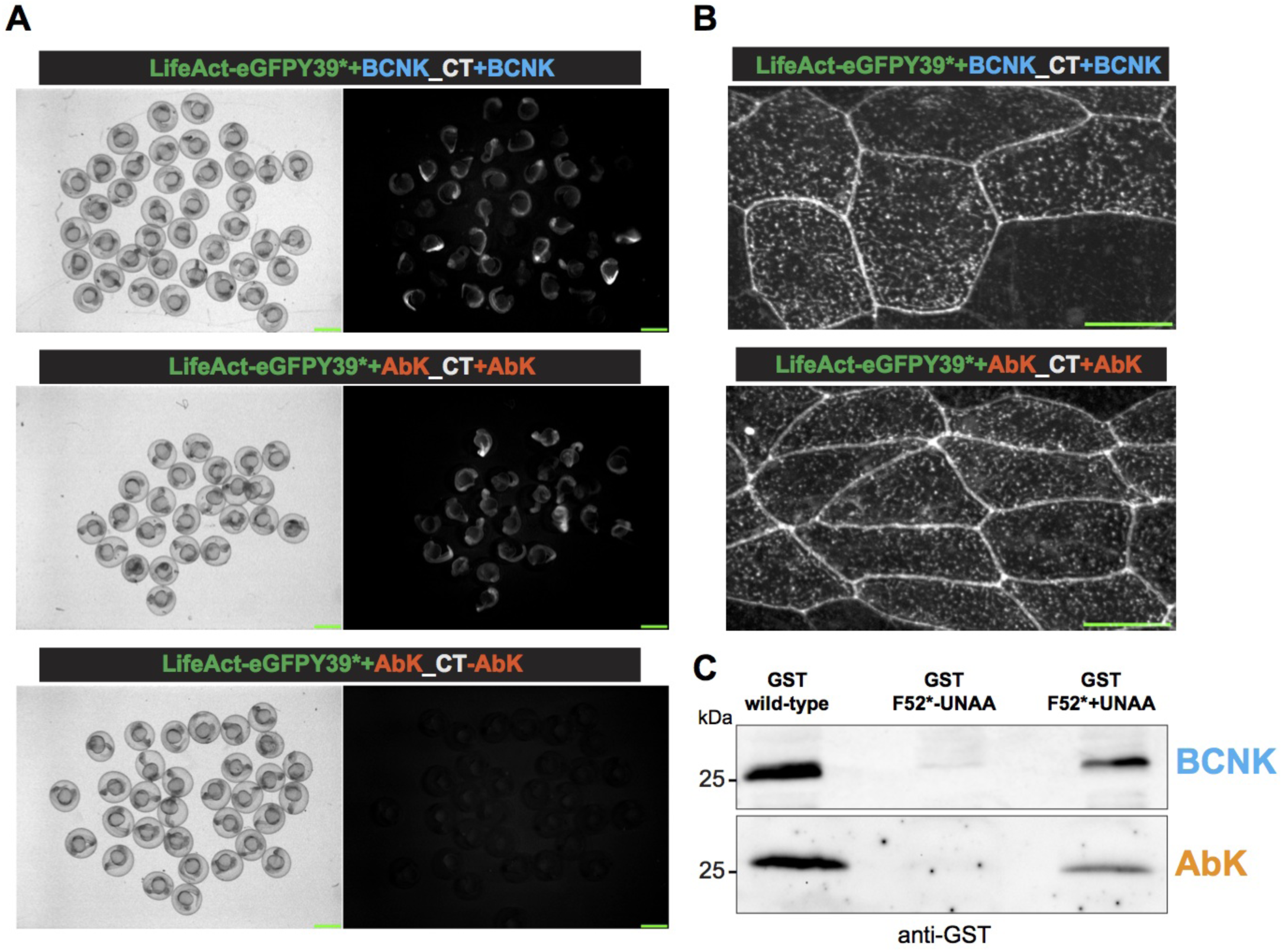
Site-specific incorporation of Bicylononyne-lysine and Diazirine-lysine in zebrafish embryos. (A). Shown are fields with multiple examples each of zebrafish embryos injected with mRNA(s) for LifeAct-eGFP Y39* with the orthogonal tRNA^Pyl-CUA^-mRNA aaRS for AzK/BCNK with (top panel) BCNK. Also shown are fields with multiple examples each of zebrafish embryos injected with mRNA(s) for LifeAct-eGFP Y39* with the orthogonal tRNA^Pyl-CUA^ -mRNA aaRS for AbK, with (middle panel) or without (bottom panel) AbK. (B). Localization of LifeAct-eGFP Y39BCNK (top panel) and LifeAct-eGFP Y39AbK to cell-cell junctions and cortical speckles. (C). Shown in the top panel is a IP-Western blot from zebrafish embryos injected with mRNA(s) for GST (left lane) or GFT F52* with the orthogonal tRNA^Pyl-CUA^ -aaRS pair for AzK/BCNK, without (middle lane) or with (right lane) BCNK. The bottom panel shows a similar experiment showing incorporation of AbK into zebrafish embryos using the appropriate amino-acid and tRNA^Pyl-CUA^ – aaRS cocktail for AbK. Immunoprecipitations and western blots were performed with antibodies against GST.

In summary, we present evidence for site-specific incorporation of three unnatural amino acids through genetic code expansion into two different reporter proteins in zebrafish embryos. These amino-acids can be used in photo-crosslinking and / or click chemistry experiments. While we have clearly demonstrated incorporation of an UNAA into GFP through proteomics experiments, further developments are needed to make this method useful in a large number of biological studies. First, we have been unable to carry out photo-crosslinking under conditions in which we have successfully carried out photo-crosslinking in bacterial, yeast and mammalian cells. Second, the UNAAs need to be injected into embryos, since in our conditions, even after removal of the chorion, they do not pass cell membranes. It is noteworthy that Deiters and colleagues have also injected lysine-derived UNAAs into zebrafish embryos [12]. Therefore, conditions that facilitate membrane permeability of UNAAs into zebrafish embryos need to be investigated. Finally, transgenic lines expressing the orthogonal tRNA-aaRS need to be created, such that this powerful technology can be used in free feeding adult fish. Notwithstanding these limitations, the methods we report provides a platform for further development of this powerful technology to enable precise engineering of proteins in zebrafish embryos. In addition to the traditional uses of this system, the fact that expression of marker proteins is only achieved in the presence of the UNAA suggests that the method we describe can also be used for inducible expression of proteins in zebrafish.

## Materials and Methods

### Un-natural amino acids (UNAAs)

AzK (N3-Lysine (N6-((2-azidoethoxy) carbonyl)-L-Lysine, SC-8027) and Exo-BCNK (Bicyclo [6.1.0] nonyne – Lysine, SC-8016) were purchased from Sirius fine chemicals SiChem GmbH. AbK (N6-[[2-(3-Methyl-3H-diazirin-3-yl) ethoxy] carbonyl]-L-lysine, 5113) was purchased from Tocris. 200 mM stock solutions of each UNAA was prepared by dissolving them in 85% 0.2M NaOH and 15% DMSO.

### Plasmid construction

The two different tRNA synthetases 1) the double mutant of wild-type *Methanosarcina mazei* PylRS (Y360A and Y384F) for incorporating AzK and BCNK and 2) the mutant *Methanosarcina barkeri PylRS (*L274M, C313A, Y349F) for integrating AbK were cloned into the pCS2+ vector. Wild type LifeAct-eGFP and GST were cloned into the pCS2+ vector. The TAG codons were introduced at Y39 position of eGFP and at Y52 position of GST respectively. All cloning procedures were performed using the in-fusion cloning kit (Clontech).

### In vitro transcription

The coding sequences cloned into the pCS2+ vector were linearized by NotI digestion. 500ng of the linearized product was used as a template in a 20µl reaction to synthesize the corresponding mRNA using mMESSAGE mMACHINE SP6 Transcription Kit (AM1340) which was further purified by phenol:choloform:isoamyl alcohol extraction.

The DNA template for synthesizing amber suppressor pyrrolysine tRNA (PylT) of Methonosarcina species were ordered from sigma as oligonucleotides preceded by the T7 promoter.

The sequences of the templates are as follows

PylT forward: 5’-ATTCGTAATACGACTCACTATAGGAAACCTGATCATGTAGATCGAATGGACTCTAAATCCGTTCAGCCGGGTTAGATTCCCGGGGTTTCCGCCA-3’ PylT Reverse: 5’-TGGCGGAAACCCCGGGAATCTAACCCGGCTGAACGGATTTAGAGTCCATTCGATCTACATGATCAGGTTTCCTATAGTGAGTCGTATTACGAAT-3’

The above-mentioned complementary oligonucleotides were annealed (annealing buffer: 10 mM Tris pH 8.0, 50 mM NaCl, 1 mM EDTA) and 1µg of the annealed oligonucleotides were used as template for *in vitro* transcription following the mMESSAGE mMACHINE™ T7 Transcription Kit protocol.

### Microinjection of Zebrafish embryos

Wild type embryos were obtained by natural mating using standard procedures in accordance with institutional animal care regulations at the University of Warwick. One–cell stage zebrafish embryos were injected with 2nl from the total 1.5µl injection mixture (0.25µl of 200 ng/µl LifeAct-eGFP/GST mRNA, 0.75µl of 200 ng/µl tRNA synthetase mRNA, 0.25µl of 4000 ng/µl of tRNA, 0.25µl of 100 mM UNAA). The injected embryos were collected 24 hours after fertilization for biochemical / cell biological analyses.

### Imaging

Embryos were incubated at 28.0°C overnight after injection. Clutches of chorionated embryos were imaged at approximately 24 hours post fertilisation on plastic petri dishes using a SMZ18 stereo microscope equipped with a Teledyne Phometrics CoolSnap HQ2 CCD camera. A Nikon P2-SHR Plan Apo 1×/0.15 NA objective lens was used to obtain a final pixel size of 8.85 µm/pixel. A Nikon Intenslight C-HGFI illuminator was used for excitation. Image acquisition was automated using Fiji micro-manager (Edelstein et al., 2010). Embryos were thence dechorionated and embedded in 0.7 % low melting agarose on No. 0 coverslips (MatTek 35 mm uncoated dishes). The dorsal anterior was imaged at approximately 24 hours post fertilisation using an Andor Revolution XD confocal system, assembled on a Nikon Eclipse Ti inverted microscope with a spinning disc confocal Yokogawa CSU-X1 unit, and an Andor iXon Ultra EMCCD camera. A Nikon Fluor 40×/1.30 NA objective lens was used to obtain a final pixel size of 0.2 µm/pixel. All images were acquired with a Z-step size of 0.5 µm. A 561 nm laser line was used for excitation. Image acquisition was automated using Andor IQ3 software. A maximum intensity projection of the optical stack was performed using Fiji (Schindelin et al., 2012).

### Western blotting and Immunoprecipitation

One–cell stage zebrafish embryos were injected with the appropriate cocktail (Orthogonal tRNA synthetase, tRNA and GST^F52^TAG or LifeAct^Y39^TAG-eGFP RNA) and UNAAs (AzK, AbK and BCNK). Embryos were collected after 24h and lysates are prepared using standard RIPA buffer (50 mM Tris (pH 8.0); 150 mM NaCl; 1% NP-40; 0.5% deoxycholate; 0.1% SDS and 1X cocktail protease inhibitors) and incubated with the buffer on ice for 15 minutes, and further clarified at 14,000 RPM for 15-20 minutes at 4°C. Clarified lysates were heated for 5 min at 95°C with 4x sample loading buffer and loaded on 12% SDS-PAGE gels, transferred to nitrocellulose membrane (GE Healthcare), and immunoblotting was performed using anti-GFP-HRP (sc-9996 HRP) and anti-GST-HRP (sc-138 HRP).

Lysates from embryos (>800) were prepared as mentioned above for the immunoprecipitation experiments. Clarified lysates were incubated with pre-washed GFP-Trap (chromotek) in RIPA buffer for 12-16hr at 4°C. GFP-Trap beads were washed with RIPA buffer 5-6 times prior loading onto 12% SDS-PAGE gels for mass spectrometry.

### Sample preparation

Sample preparation: Samples from zebrafish embryo extracts were run in 12 % SDS-PAGE minigels until the dye front was 1 cm from the bottom. The gels were washed with deionised water three times (5 minutes each) and stained with Coomassie blue (SimplyBlueStain, Invitrogen) overnight and destained with deionised water for 4-6 hrs. LifeAct-eGFP carrying unnatural amino acid (AzK) band (around 27-29 KDa) in the gel was cut into cubes of ∼1 mm^3^ for in gel tryptic digestion [19] prior to MS analysis.

### NanoLC-ESI-MS/MS analysis and UNAA identification

Reversed phase chromatography was used to separate tryptic peptides prior to mass spectrometric analysis. Two columns were utilised, an Acclaim PepMap µ-precolumn cartridge 300 µm i.d. × 5 mm 5 µm 100 Å and an Acclaim PepMap RSLC 75 µm × 25 cm 2 µm 100 Å (Thermo Scientific). The columns were installed on an Ultimate 3000 RSLCnano system (Dionex). Mobile phase buffer A was composed of 0.1% formic acid in water and mobile phase B 0.1 % formic acid in acetonitrile. Samples were loaded onto the µ-precolumn equilibrated in 2% aqueous acetonitrile containing 0.1% trifluoroacetic acid for 5 min at 10 µL min-1 after which peptides were eluted onto the analytical column at 250 nL min-1 by increasing the mobile phase B concentration from 4% B to 25% over 37 min, then to 35% B over 10 min, and to 90% B over 3 min, followed by a 10 min re-equilibration at 4% B [20].

Eluting peptides were converted to gas-phase ions by means of electrospray ionization and analysed on a Thermo Orbitrap Fusion (Thermo Scientific). Survey scans of peptide precursors from 375 to 1575 m/z were performed at 120K resolution (at 200 m/z) with a 2 × 105 ion count target. Tandem MS was performed by isolation at 1.2 Th using the quadrupole, HCD fragmentation with normalized collision energy of 33, and rapid scan MS analysis in the ion trap. The MS2 ion count target was set to 1×104 and the max injection time was 200 ms. Precursors with charge state 2–6 were selected and sampled for MS2. The dynamic exclusion duration was set to 25 s with a 10 ppm tolerance around the selected precursor and its isotopes. Monoisotopic precursor selection was turned on. The instrument was run in top speed mode with 2 s cycles.

## Data Analysis

The raw data were searched using MaxQuant [21] (version 1.6.2.6) against the sequence of the construct, Uniprot Danio rerio reference proteome (www.uniprot.org/proteomes/UP000000437) and the MaxQuant common contaminant database. For the database search, peptides were generated from a tryptic digestion with up to two missed cleavages, carbamidomethylation of cysteines as fixed modifications and variable modifications used were oxidation of methionine, acetylation of the protein N-terminus, Azido-Lysine (+C3H3O2N3).

## Author Contributions

JS initiated the zebrafish experiments, generated the tRNA, aaRS, marker protein mRNA constructs. JS and SP carried out the first successful genetic code expansion experiments in the MKB group. SP carried out all biochemical studies and assembled the manuscript data figures. STC performed all zebrafish-imaging experiments. JS, SP, and STC also helped with preparation of the manuscript. ZA provided expert technical assistance. SP, ARB and AMEJ performed the proteomic analysis. KS trained JS and supervised the initial zebrafish experiments. MKB conceived the study, secured funding, supervised the project and wrote the first draft of the manuscript. All authors edited and approved the manuscript.

## Funding

This work was supported by Wellcome Trust Senior Investigator Award (WT101885MA), a Royal Society Wolfson merit award (WM130042) and an European Research Council Advanced Grant (ERC-2014-ADG N° 671083) to MKB. KS is supported by the BBSRC. During the early stages of the project, STC was funded by a studentship from the Engineering and Physical Sciences Research Council (EP/F500378/1).

### Acknowledgements

Special thanks are due to Dr. Kensaku Sakamoto (RIKEN, Yokohama, Japan) for generously providing plasmids used in this study as well as hosting JS in his laboratory to learn general genetic code expansion methods. We thank all members of the MKB laboratory, especially Ms. Paola Zambon and Dr. Tomoyuki Hatano, for discussion and feedback on experiments. We thank the University of Warwick Biological Services Unit for the expert housing and care of zebrafish used in this study.

